# Uracil/H^+^ symport by the FurE transporter challenges the rocking-bundle mechanism of transport in APC transporters

**DOI:** 10.1101/2022.03.28.486045

**Authors:** Iliana Zantza, Georgia F. Papadaki, Stefano Raniolo, Yiannis Pyrris, George Lambrinidis, Vittorio Limongelli, George Diallinas, Emmanuel Mikros

**Affiliations:** Department of Pharmacy, National and Kapodistrian University of Athens, Panepistimiopolis, Athens, 15771, Greece; Department of Biology, National and Kapodistrian University of Athens, Panepistimiopolis, Athens, 15781, Greece; Faculty of Biomedical Sciences, Euler Institute, Università della Svizzera italiana (USI), Lugano, 6900, Switzerland; Department of Pharmacy, University of Naples “Federico II”, Naples, 80131, Italy; Institute of Molecular Biology and Biotechnology, Foundation for Research and Technology, Heraklion, 70013, Greece; Athena Research and Innovation Center in Information Communication & Knowledge Technologies, Marousi, 15125, Greece

**Keywords:** Aspergillus nidulans, NCS1, fungal, specificity, structure-function

## Abstract

Transporters mediate the uptake of solutes, metabolites and drugs across the cell membrane. The eukaryotic FurE nucleobase/H^+^ symporter of *Aspergillus nidulans* has been used as a model protein to address structure-function relationships in the APC transporter superfamily, members of which are characterized by the LeuT-fold and seem to operate by the so-called ‘rocking-bundle’ mechanism. In this study, we reveal the binding mode, translocation and release pathway of uracil/H^+^ by FurE, using path collective variable, funnel metadynamics and rationally designed mutational analysis. Our study reveals a step-wise, induced-fit, mechanism of ordered sequential transport of proton and uracil, which in turn suggests that the FurE symporter, and probably structurally similar transporters, functions as a multi-step gated pore, rather than employing ‘rocking’ of compact domains, as generally proposed for APC transporters. In addition, our work further supports the emerging concept that specific elements of cytosolic terminal regions of transporters might be functionally important.

## Introduction

Secondary active transporters are transmembrane proteins that mediate the transport of nutrients, metabolites and drugs in or out of cells. They select and translocate their substrates using the energy provided by the electrochemical gradient of the membrane, via a mechanism that involves the symport or antiport of mostly Na^+^/H^+^ cations with other solutes. Structural studies revealed that although secondary active transporters may be structurally, functionally or evolutionary distinct, they share common folds, which are related to specific protein conformational changes associated with the transport cycle. The general model for the transport mechanism is known as the ‘alternating-access model’, where the transporter accepts or releases the substrate at one side of the cell membrane by changing conformations from an *outward-open* (OO) state facing the extracellular environment to an *inward-open* (IO) state facing the cytosol.^1–5^ Depending on the folding and specific conformational rearrangements of the transporter, three major mechanisms have been proposed, namely the rocker-switch, the rocking-bundle and the sliding-elevator.^5–9^

Important structural and functional information about the rocking-bundle mechanism, which characterizes one of two largest transporter families, the so-called Amino Acid-Polyamine-Organocation (APC) superfamily, rise from seminal studies on the bacterial transporter LeuT, specific for leucine and alanine.^3,5,10^ LeuT adopts the 5+5 helical inverted repeat (5HIRT), formed by the first 10 transmembrane helices whose structural elements and conformational changes determine substrate recognition and transport. In total, LeuT and most APC transporters possess twelve transmembrane α-helical segments (TMSs), however the role of TMS11 and TMS12 is not yet clarified. The rocking-bundle model assumes that translocation of the substrate following the OO-to-IO conformational change is facilitated by the relative motion between two motifs, the so-called ‘hash’/scaffold domain (TMS3, TMS4, TMS8, TMS9) and the ‘bundle’/core domain (TMS1, TMS2, TMS6, TMS7), with TMS5 and TMS10 functioning as gates. It has been suggested that substrate binding in the OO conformation is assisted by the simultaneous binding of a positive charge ion (Na^+^ or H^+^), which elicits the conformational change of the protein towards the IO conformation. This mechanism of substrate translocation has been supported by studies on the eukaryotic dopamine (DAT)^11^ and serotonin (SERT)^12^ transporters (neurotransmitter/sodium symporter family-NSS), and a number of mostly prokaryotic transporters.^13–22^

Although all transporters conforming to the 5+5 APC structure share the same ‘bundle-hash’ fold, topological differences have been found during the transition from the OO to the IO state. LeuT and MhsT crystal structures suggest that the ‘bundle’ domain (TMS1, TMS2, TMS6, TMS7) undergoes significant conformational changes during the OO/IO transition, pivoting around the ‘hash’ domain (TMS3, TMS4, TMS8, TMS9), while there are two additional rearrangements functioning as opening-closing gates. In LeuT, specifically, the displacement of TMS1b, TMS6a acts as an extracellular gate, along with a 45-degree kink of the TMS1a followed by a local unwinding of TMS5, which functions as the intracellular gate. In contrast, the Mhp1 transporter transits from the outward- to the inward-state by rocking a mobile ‘hash’ motif around the ‘bundle’ domain, which also promotes TMS10 to move towards TMS1b and TMS6a to pack the substrate in the occluded conformation. Additionally, a flexible TMS5 bending, rather than movements in TMS1a of LeuT, opens the inward facing cavity and facilitates substrate release, thus functioning as the inner gate.

Several fungal members of the nucleobase cation symporter 1 (NCS1) family, which are structurally related to the APC superfamily, have been extensively studied by Diallinas and co-workers, unveiling important information about regulation of expression, subcellular trafficking and turnover, transport kinetics, and substrate specificity.^23–29^ Transporters of this family function as H^+^ symporters selective for uracil, cytosine, allantoin, uridine, thiamine or nicotinamide riboside and secondarily for uric acid and xanthine.^23,24,30^ In previous studies, we have modeled several NCS1 transporters of *Aspergillus nidulans* using the prokaryotic Mhp1 benzyl-hydantoin/Na^+^ transporter as a structural template, and assessed structure-function relationships via extensive mutational analyses. From these studies, we defined the substrate binding site and revealed the important role of the cytosolic N-and C-terminal segments in regulating endocytic turnover, transport kinetics and surprisingly substrate specificity,^25–29^ the importance of N-terminus in transporter function has been also proved in the case of hSERT.^31^

Here, we sought to describe the functional conformational changes associated with the transport activity of the most extensively studied fungal NCS1 member, namely the FurE uracil/uric acid/allantoin transporter, from the OO to the IO state. To this end, we employed metadynamics calculations^32^ and additional mutational analyses, rationally designed to assess our *in silico* findings. Overall, we were able to characterize the large-scale conformational changes of FurE from the OO to the IO state, including several intermediate states, elucidating the role and the internalization order of both substrate (uracil) and H^+^ (in the form of H_3_O^+^ and their binding modes, thus providing a comprehensive novel picture that challenges aspects of the rigid-domain rocking mechanism of APC transporters.

## Results

### FurE 3D structure

The FurE structure in three different conformational states, Outward Open (OO), Occluded (Occ) and Inward Open (IO), was built through homology modeling using the corresponding Mhp1 crystal structures (**Figure 1 and Figure S1**).^20–22^ Upon visual inspection of the structures, it emerges that interactions between residues are expected to be crucial for the structure and function of the transporter. For example, R123 (TMS3) can form a salt bridge with D261 (TMS6), mimicking the interaction observed in Mhp1 between K110 (TMS3) and D229 (TMS6) (**Figure S2A**). Another important interaction is between E51 at the edge of TMS1a and K199 of TMS5 (**Figure S2B**). Interestingly, in both OO and Occ cases, the K199 side-chain amino group is situated in the position of the co-crystallized Na^+^ cation in the Occ conformation of Mhp1 and very close to that of the second Na^+^ (Na2) present in the equivalent structure of LeuT (**Figure S2C**). Additionally K252 (TMS6), which has been shown to affect substrate specificity,^28^ might also form a second salt bridge with E51 (**Figure S2B**). Finally, the cytoplasmically located N-terminal D28 appears to interact with K188 (TMS5) in OO, as also reported by Papadaki *et al*., ^29^ and with R264 in Occ (**Figure S2A**). Apart from the aforementioned ‘static’ salt bridges, additional interactions, possibly involved in the function of the outer gate could be between the Q59, T63, S64 side chains (TMS1b) and F385, S386 (TMS10) (**Figure S2D**), while hydrophobic interactions involving W39 might control the inner gate.

**Figure 1.**
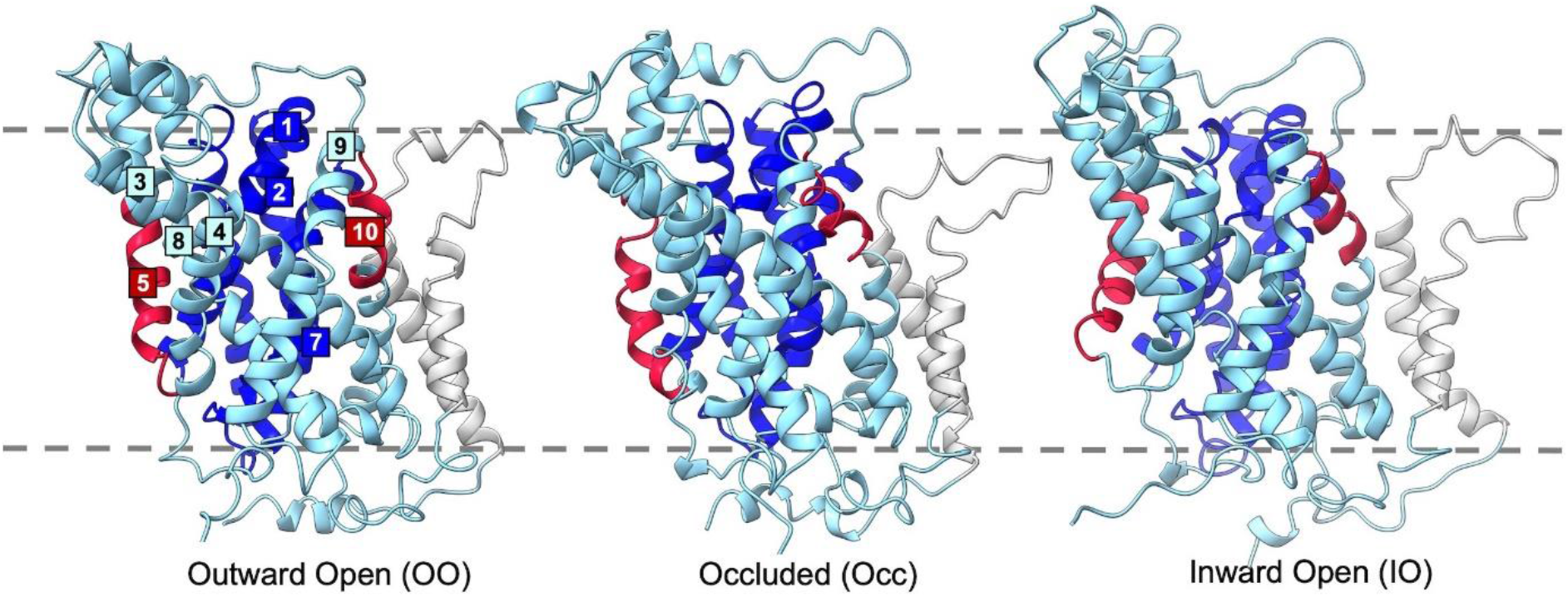
Model structures of FurE. The three homology models, Outward Open (OO), Occluded (Occ), Inward Open (IO), based on corresponding Mhp1 template crystal structures are shown in side view (orientation parallel to the membrane lipids). The ‘bundle’ helices are colored blue, the ‘hash’ helices are colored cyan, the outer and inner gates are colored red, and the TMS11, and TMS12 are grey. The yellow dashed lines represent the membrane plane.

### Mutational analysis confirms the crucial role of specific residues in FurE transport function

The FurE structural models highlight two salt bridges, R123-D261 and E51-K199, and a polar interaction between Q59 and S384 or S386 as crucial in the conformational transitions of FurE. Additional residues predicted to be related to conformational changes were W39, T63, S64, R193, F196, R264, N347 and F385. In order to support these predictions, we performed respective Ala substitutions. Other residues predicted to be important for transport activity, such as the interaction of D28 with K188, and the critical role of K252 in substrate binding and specificity, have been previously studied by analogous Ala substitutions.^29^ Mutated versions of FurE, C-terminally tagged with GFP, were analyzed in an *A. nidulans* (Δ7) strain that genetically lacks all major nucleobase-related transporters, as previously described^28,29^. **Figure 2A** (upper left panel) summarizes growth phenotypes of mutants and control strains. As expected, the positive control strain expressing wild-type FurE grows on allantoin and uric acid and is sensitive to 5-fluorouracil (5-FU), whereas the negative control strain not expressing FurE shows a N starvation growth phenotype and is resistant to 5-FU. Ala substitutions in residues predicted to form the two major salt bridges (R123-D261 and E51-K199) scored as loss-of-function mutations, reflected in abolishment or dramatic reduction of growth on allantoin or uric acid and relatively increased resistance to 5-FU, mostly evident in R123A and D261A. Similar dramatic loss of FurE transport activity was obtained in R264A and F385A mutants, while Q59A and S386A FurE versions seem to have lost their transport activity for uric acid or 5-FU, but conserved some capacity for allantoin transport. Thus, the mutational analysis confirms the essential functional role of the interactions between R123-D261, E51-K199 and Q59-S385-S386, as well as the importance of R264, which is predicted to interact with N-terminal D28. The mutational analysis also revealed an important role of W39, as its substitution led to loss of FurE-mediated uric acid and allantoin transporter, although 5-FU transport to this drug is retained. Our findings further showed that Ala substitution of T63, S64, R193 or F196 have moderate negative effects on FurE apparent activity, reflected in reduction of growth on uric acid and some increase in 5-FU resistance (e.g., F196A), whereas residues N347 and S384 proved non-essential for FurE activity.

**Figure 2:**
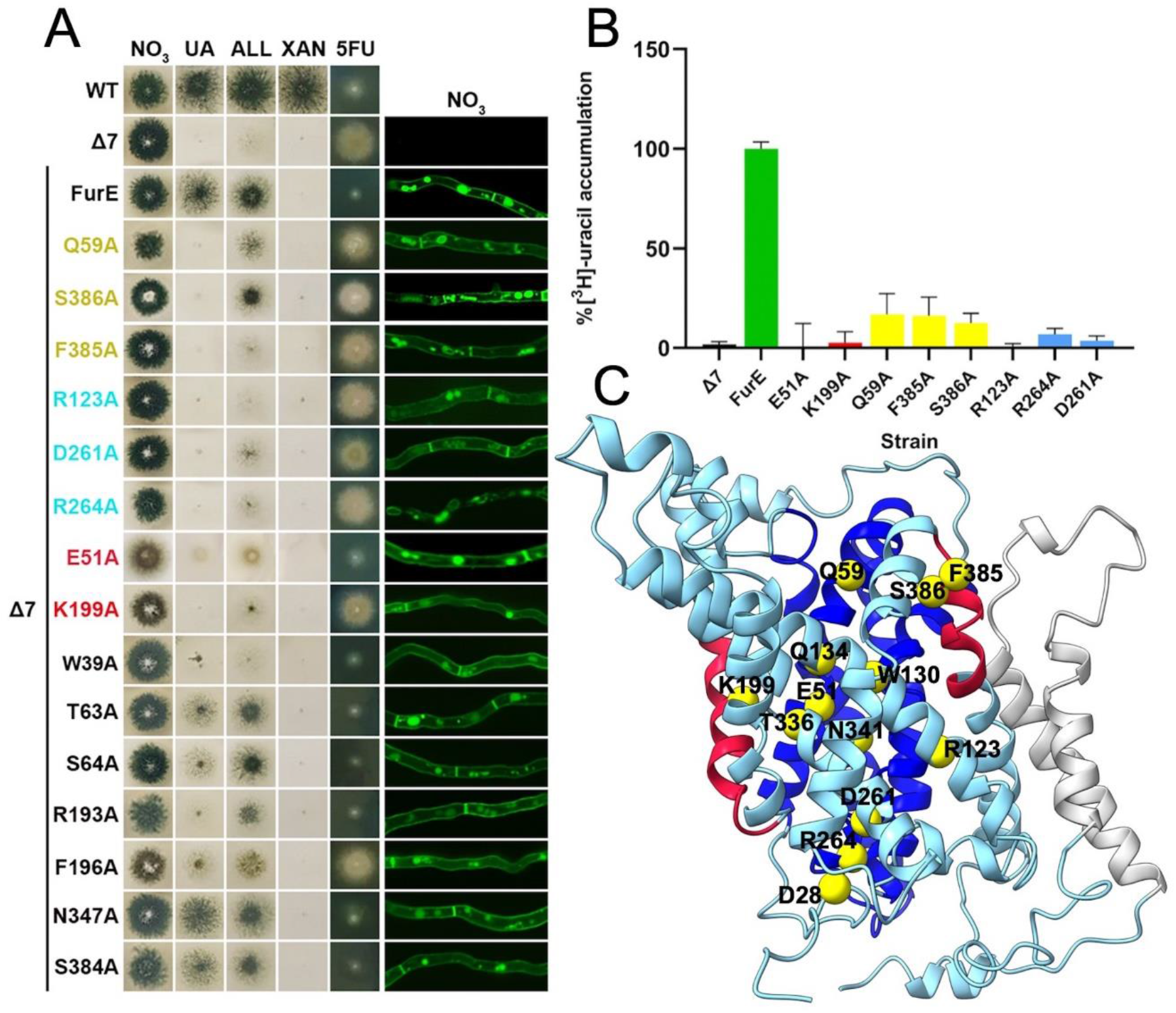
Functional analysis of FurE mutants. **(A)** Growth tests of isogenic strains expressing distinct FurE mutant versions in a Δ7 genetic background (i.e. genetically lacking other nucleobase-related transporters), compared to a positive (FurE) and a negative (Δ7) control strain (for strain details see materials and methods). NO3, UA, ALL, Xan denote MM supplemented with nitrate, uric acid, allantoin or xanthine as sole N source. 5FU is MM+NO3 supplemented with 5-FU. WT denotes a standard *A. nidulans* wild-type strain expressing all major nucleobase transporters. *In vivo* epifluorescence microscopy of the same strains is shown in the right panel. All FurE mutants are functionally tagged with GFP. Notice that all FurE mutant versions, except R264A, exhibit normal (i.e. wt FurE-like) plasma membrane localization and vacuolar turnover. R264A is trapped in the perinuclear ER rings, typical of misfolded versions of FurE or other transporters (for details see Materials and methods) (B) Direct uptake assays of selected FurE mutants, using 0.2 μM [3H]-radiolabeled uracil. The figure shows relative % initial uptake rates (1 min) of mutants, when wild-type FurE transport is taken as 100%, performed with 107 germinated conidiospores, as described by Krypotou and Diallinas, 2014 (for details see Materials and methods)

Epifluorescence microscopic analysis, shown in the right panel of **Figure 2A,** confirmed that mutational disruption of the major interactions tested (R123-D261, E51-K199 and Q59-F385-S386) did not affect the normal PM localization and stability of FurE, which confirms that the associated growth defects in specific mutants reflect defects in FurE transport activity *per se*, rather than an effect on protein folding or subcellular trafficking. Direct transport assays–showed that FurE-mediated radiolabeled uracil transport was abolished in the respective mutants (**Figure 2B).** Noticeably, only in the case of R264A mutant the apparent loss-of-function proved to be the result of abolishment of trafficking to the PM, due to ER-retention of FurE. In conclusion, mutations of residues proposed, via homology modeling and initial MDs, to be functionally important validated the structural models constructed.

### The binding mode of hydronium

Contrastingly to Mhp1, which is a Na^+^-driven NCS1 symporter, all characterized fungal NCS1 transporters function via proton (H^+^) symport. Nevertheless, proton interactions are not elucidated for none of them, including FurE. In a first step we aspire to determine possible interactions implying the proton and residues located towards the outer gate of the transporter. In this aspect we investigated the binding of a hydronium molecule (H_3_O^+^) to FurE by employing Funnel-Metadynamics (FM), developed by our group and widely used to study ligand-protein systems.^33^ During the FM calculations the whole binding pathway was simulated and all possible binding sites were energetically evaluated (**Figure S3A**). The preferential binding site of hydronium was identified as the lowest energy state in the Binding Free Energy Surface (BFES) (**Figure 3A**) and proved to be the same site identified for Na^+^ in both Mhp1 and LeuT sodium co-crystallized structures. This site, located at the interface of TMSs 1 and 8, involves E51 (TMS1b) and T336 (TMS8), the later residue conserved also in LeuT and Mhp1 (**Figure 3B)**. The structural stability of the binding complex FurE/H_3_O^+^ was further assessed by a 150 ns MD simulation.

**Figure 3:**
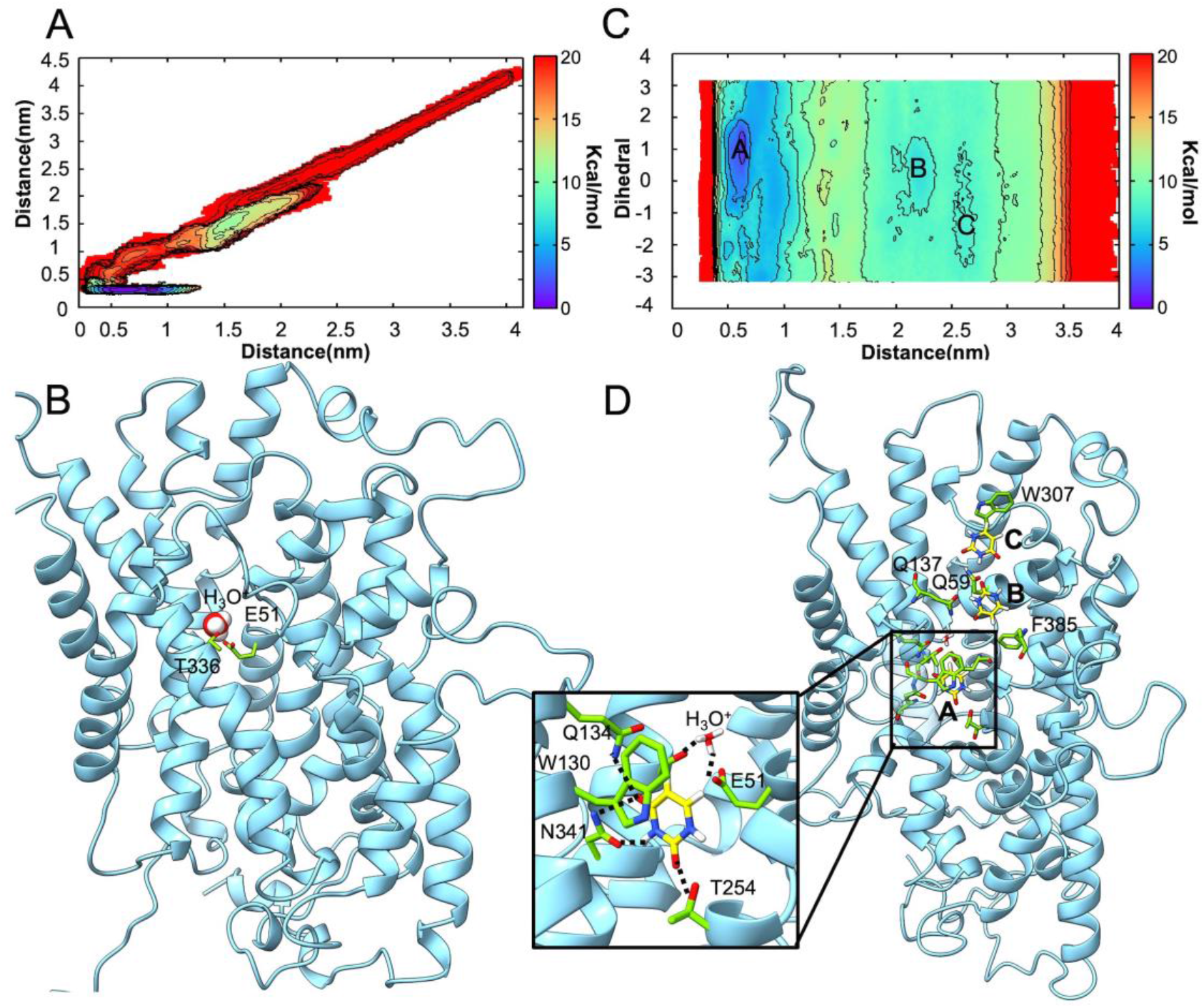
Binding of H_3_O^+^ and uracil as simulated by Funnel Metadynamics. (A) The BFES of H_3_O^+^ binding process. Contour lines are shown every 2 kcal/mol. (B) The binding mode of H_3_O^+^ cation in FurE transporter as derived from the global energy minimum in the FES. (C) The BFES of uracil binding process in FurE transporter. (D) The intermediate states (local minima in the BFES) of uracil entering FurE transporter and the binding mode in the binding site (inset) as derived from the BFES in C.

### The binding mode of uracil

Next, we simulated the binding process of uracil (K_m_ = 1 mM) to its putative binding site in FurE using FM. The putative binding site was confined between TMS1, TMS3, TMS6 and TMS8 as suggested by previous mutagenesis data, as well as structural studies on Mhp1 and other NCS1 transporters.^26^ As performed in the case of H_3_O^+^, we simulated the binding process of uracil from its fully solvated state to the binding site in the Occ state, using the uracil distance to the binding site as CV. Uracil starting structure was generated by docking calculations, which does not affect the final result since FM calculations explore all the possible binding poses. To ensure a wide area sampling around the binding site we have set a large cone section in the FM simulation (**Figure S3B**). H_3_O^+^ was also included at the binding mode previously identified according to the existing literature.^19,22^

Derived from the global minimum of the BFES (**Figure 3C**), the selected model of uracil binding mode (**Figure 3D**) was found to be remarkably similar to that of (5S)-5-benzylimidazolidine-2,4-dione (hydantoin analogue) in the Mhp1 crystal structure (**Figure S4**).^22^ In more detail, we observed H-bond interactions between T254 (TMS6) and uracil C2=O, N341 (TMS8) and uracil C4=O and N3, and π-π stacking interactions between W130 (TMS3) and uracil. Two additional minima were found at higher energy values that represent probable intermediate binding poses of the ligand along its path to the final binding site. Uracil appears first to interact with Q59 (TMS1b), via a bidentate interaction with N3 and C2=O, and a π-π stacking with W307 (L7 loop) (**Figure 3C**). Subsequently, it moves lower in the FurE binding cavity, where it interacts with Q137 (TMS3) via a bidentate bond involving C4=O and N3 (**Figure 3D**). Finally, uracil and W130 both interacted with F385 (TMS10) through π-π and T-shaped stacking interactions.

### The conformational transition of FurE from OO to IO

To thoroughly describe the large-scale conformational transition of FurE, and the relative order with which hydronium and uracil are transported, we employed a dimensionality reduction approach, called path collective variables (PCVs).^34^ In this case, the aforementioned transitions can be discretized by providing a set of frames describing the required movement (see Methods). These frames include the positions of key atoms from the beginning to the end of the conformational change, allowing us to track the transition stage during the simulation and also accelerate its sampling through Metadynamics. The whole transition of FurE from OO to IO was investigated through two set of simulations, the first describing the OO-to-Occ transition and the second the Occ-to-IO. For each of them, four systems were investigated considering all possible combinations of ligand stoichiometry: i) FurE -H_3_O^+^ - uracil (*apo*); ii) FurE + H_3_O^+^ - uracil; iii) FurE - H_3_O^+^ + uracil; iv) FurE + H_3_O^+^ + uracil (the total simulation time for each metadynamics is shown in **Table S1**). In the simulations where hydronium and uracil are present, they occupy the binding mode previously identified. FurE structures representing the global minimum at the calculated FES were extracted and clustered. The centroid structure of the most populated cluster was selected and subjected to a 100ns standard MD simulation in order to assess its stability. The interactions between the most important residues were monitored within the extracted structures and statistics are shown in **Figure 4A**.

**Figure 4.**
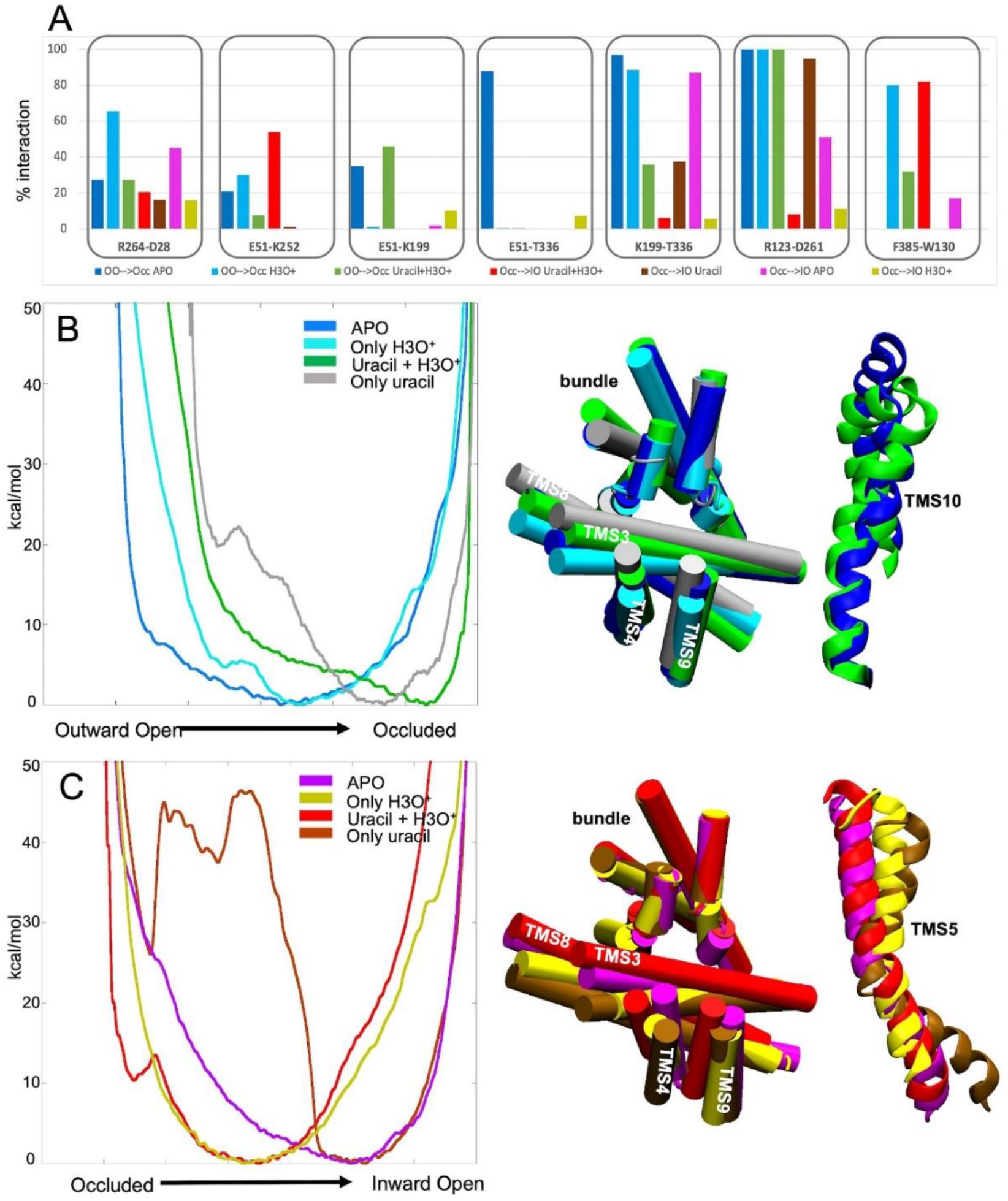
FurE structural alterations, residue interactions and Free Energy Surface plots during transport conformational changes. (A) Side chain interactions of important residues have been monitored in all structures collected in each FES global minimum and are represented as percentage over the ensemble of the structures. (B) The FESs of the OO-to-Occ transition using different stoichiometry of ligands bound to the transporter (colour code: protein in the *apo* form in blue, complexed only with H_3_O^+^ in cyan, complexed with both uracil and H_3_O^+^ in green, complex only with uracil in grey). Each tick in the x axis represents one unit. (C) The FESs of the Occ-to-IO transition are shown using different stoichiometry of ligands bound to the transporter. The system containing both uracil and H_3_O^+^ is represented in red, the system containing only uracil is represented in brown, the *apo* form is represented in magenta, the system containing only H_3_O^+^ is represented in yellow.

### The OO-to-Occ transition

#### i) Apo state (FurE - H_3_O^+^ - uracil)

The FES shows one single, wide energy minimum between OO and Occ (**Figure 4B**) indicating a relative conformational flexibility of FurE when no ligand is present, confirmed also by standard MD simulations (**Figure S5**). Notably, in the energy minimum structures the initial part of TMS10 is positioned much closer to TMS1a with respect to the starting OO state. The relative orientation between ‘hash’ and ‘bundle’ motives remained very similar to Mhp1.The most stable interactions in the *apo* state are engaged by K199-T336, E51-T336 and R123-D261, while E51-K199 and R264-D28 interact at a minor extent (**Figure 4A**). Additionally, S386 and Q59 can form a water bridge (**Figure S6**), while Q134 interacts with T336 through a water molecule (**Figure S7A**).

#### ii) Hydronium bound (FurE + H_3_O^+^ - uracil)

When H_3_O^+^ cation is bound to FurE, the FES is rather similar to that of the *apo* form, albeit the minimum is narrower indicating a reduced flexibility of the transporter and in particular of TMS10 (**Figure 4B)**. This finding suggests that the presence of H_3_O^+^ influences the free energy landscape, leading TMS10 in a position competent to bind the substrate. More specifically, H_3_O^+^ engages in a salt bridge with E51 and H-bond with T336, disrupting the bond between K199 and E51, between E51 and T336, and the water bridge between Q134 and T336. Consequently, T336 interacts only with K199 so that the H-bond between T336 and Q134 is lost, making Q134 available to interact with uracil (**Figure S7B**).

#### iii) Uracil bound (FurE - H_3_O^+^ + uracil)

When only uracil is bound to FurE, the FES minimum was located close to the Occ state (**Figure 4B**). However, in this pose the first part of TMS10 is still relatively distant to TMS1a. Unbiased MD simulations performed on this system show that uracil is not stable in the binding pocket, leaving the binding mode after 20ns, while TMS10 fluctuated between an OO and Occ state (**Figure S5**).

#### iv) Hydronium and uracil bound (FurE + H_3_O^+^ + uracil)

In the case both hydronium and uracil are bound, the lowest energy minimum represents the Occ state (**Figure 4B)**. Comparing this FurE state to the crystallized Occ state of Mhp1, minor differences are observed in TMS5, where a tilt was noted towards the IO conformation, and in TMS3 and TMS9. A network of interactions between F385, S386, F388, L389 with Q59, V60 and W130 contributes in stabilizing TMS10 in the occluded position, with a consequent motion of TMS9, which however is not observed in the Mhp1 crystal structure. Furthermore, the slight tilt of TMS5 suggests that FurE in Occ has already moved in a conformation closer to the IO state, foreshadowing a low energy barrier between the occluded and an Inward Occlude state (IOcc). Additionally, binding of uracil stabilizes Q134 through an H-bond, in a position capable of making an H-bond network with Q59 and water molecules. S386 (TMS10) can interact with Q59 either via a water molecule or directly (**Figure S6**).

Taken together, our results provide unprecedented structural insight into the OO-to-Occ transition of FurE. In detail, it is clearly shown that the presence of hydronium stabilizes the FurE conformation competent for binding the uracil and that the binding of both hydronium and uracil is necessary to lock FurE in Occ state. Our observations agrees with experimental data concerning Mhp1, where the symported Na^+^ increase substrate affinity without inducing major conformation changes,^19,35^ while in LeuT Na^+^ binding shifts the conformational equilibrium to the Occ state.^36,37^ Occ state in FurE is a very stable state as demonstrated by the low RMSD values (~1 Å) computed for the backbone Cα atoms of the transporter in unbiased MD calculations. Additionally, the break of initial bonds that stabilized TMS5 in a closed position and retained a stable ‘hash-bundle’ domain orientation (K199-E51, K199-T336, E51-T336), allow the FurE structure to move towards IO state.

### The Occ-to-IO transition

#### i) Hydronium and uracil bound (FurE + H_3_O^+^ + uracil)

When uracil and H_3_O^+^ are both bound to FurE, the transporter assumes a low energy structure that approaches the IO state, albeit not reaching it. This can be defined as the Inward Occluded (IOcc) state. Here, TMS3 is tilted, inducing TMS4 and TMS5 to assume a semi-open state. At the same time the central part of TMS8 is tilted away from TMS1a. The TMS3 motion is characterized by the break of the electrostatic interaction between D261 (TMS6) and R123 (TMS3) (**Figure 4A**), which instead interacts with T254 of the uracil binding site and uracil itself. H_3_O^+^ in a more interior position, approaches D28 in the N-terminal loop. In addition, the E51 side chain rotates following the cation motion and this results in a more stable interaction with K252 (**Figure 4A**).

#### ii) Uracil bound (FurE - H_3_O^+^ + uracil)

In this state, the FES shows a minimum close to the IO conformation. Such minimum is narrow, confined by a high-energy barrier (**Figure 4C**). This finding suggests that first H_3_O^+^ unbinds FurE, then the transporter is stabilized in a close-to-IO conformation useful for uracil release. Compared to the Mhp1 inward structure, the tilt of TMS5 is more pronounced, while TMS8 is not tilted anymore leading to a maximum distance from TMS1a. Both conformational changes elicit a rearrangement of the other helices of the “hash” motif TMS3, TMS4 and TMS9 (**Figure S8**).

#### iii) Apo state (FurE - H_3_O^+^ - uracil)

In the *apo* system, the FES shows the lowest energy minimum close to IO, in a position similar to the uracil bound state. However, here the minimum is wider, indicating a larger conformational freedom of the transporter in the *apo* state. TMS5 is rather flexible, while TMS3 is slightly bent if compared with the Occ state. As FurE assumes the *apo* state after the release of both the ligands, such conformational freedom might be instrumental to favor the reverse transition of the transporter to the outward state. The FurE flexibility was confirmed by standard MD simulations carried out on the structure of the minimum.

#### iv) Hydronium bound (FurE + H_3_O^+^ - uracil)

When only H_3_O^+^ is bound to FurE, the energy minimum structure is between Occ and IO (**Figure 4C**), with TMS5 very close to the position assumed in Occ. This hints that in the absence of uracil the protein is not able to reach the IO state.

Our calculations show that when H_3_O^+^ is still bound to the protein the conformation is stabilized in an intermediate state between Occ and IO. This suggests that the sequence of events includes first dislocation and dissociation of the H_3_O^+^, while uracil is needed in order to shift the Occ to the final IO state. Furthermore, this transition from Occ to IO is related to TMS3 and TMS8 tilting, a shift associated also with both H_3_O^+^ and substrate interactions.

### The internalization pathway of H_3_O^+^ cation

Based on our PCV calculations on the FurE Occ-to-IO transition, hydronium is the first to be released in the intracellular environment. Therefore, we investigated the unbinding of H_3_O^+^ from the transporter by means of FM simulations (**Figure S3C**). Our results show that hydronium is able to move towards the intracellular region of FurE passing through different binding modes (**Figure 5A, 5B**). First, H_3_O^+^ breaks the interactions with T336 to H-bond with S339, while maintaining the salt bridge with E51. This corresponds to minimum D in the FES reported in **Figure 5A.** Then, hydronium binds to a cleft created by F47, F262 and E51, corresponding to minimum C (**Figure 5A**). Afterwards, the interaction with E51 is lost and H_3_O^+^ binds to D28, D26 of the cytosolic N-terminal terminus^29^ and N347, corresponding to minimum B of the FES (**Figure 5A**). Finally, H_3_O^+^ reaches the lowest energy pose A, binding to D28 and D26 (**Figure 5B**), before being fully released in the cytoplasm. The motion of E51 along with the H_3_O^+^ unbinding elicit a break of E51-K199 and D28-R264 interactions (**Figure 4A**). Notably, our simulations indicate that the flexibility of the cytosolic N-terminal segment of FurE plays a major role in the release of hydronium in the cytoplasm.

**Figure 5.**
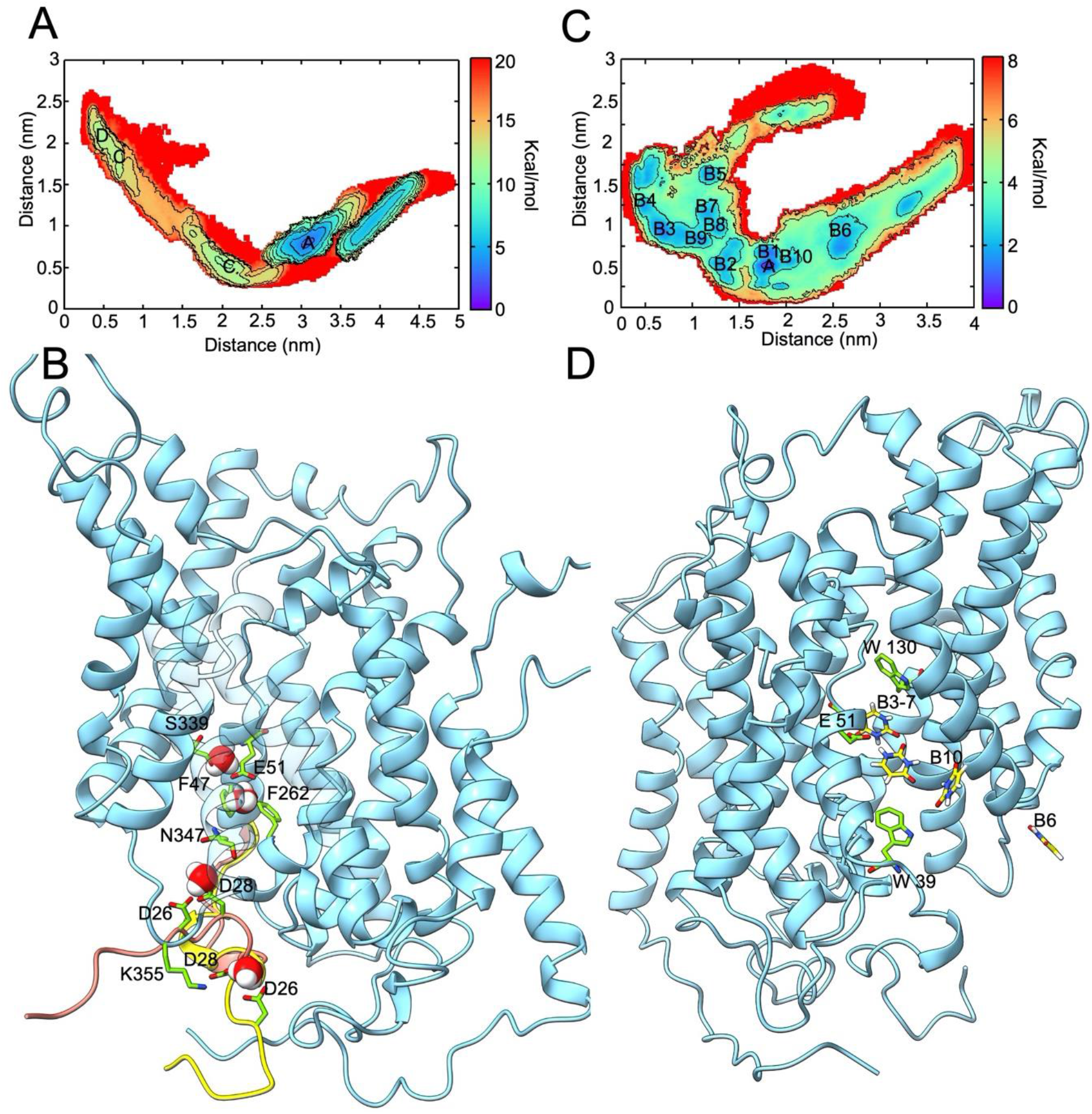
The unbinding process of H_3_O^+^ and uracil to the cytoplasm as simulated by Funnel Metadynamics. (A) The BFES of H_3_O^+^ internalization process. The separation between contours is 2 kcal/mol. (B) The binding sites of H_3_O^+^ cation in FurE transporter along the internalization pathway, as derived from the low energy states in the BFES in A. In orange is represented the cytosolic N-terminal LID when H_3_O^+^ is bound in D26, D28 and N347, while in yellow when H_3_O^+^ is released in the cytoplasm. (C) The BFES of uracil internalization process. The separation between contours is 2 kcal/mol. (D) The intermediate states of uracil internalization pathway while exiting the FurE transporter, as derived from the BFES in C.

### The internalization pathway of uracil

Once hydronium unbinds, uracil can be released in the intracellular environment. We investigated the unbinding of uracil from the FurE IO state investigating all the possible exiting pathways from the binding pocket to the TMS5 inner gate (**Figure S3D**). The FES and the ligand energetically relevant poses are represented in **Figure 5C, 5D,** respectively. During uracil unbinding three residues, W130, E51 and W39, play a major role (**Figure 5D**). W130 keeps a vertical conformation to the z axis of the membrane, thus closing *de facto* the access to the extracellular part, while E51 forms a H-bond with uracil favoring the translocation of the ligand towards the TMS5 inner gate. Finally, W39 forms π-π and T-shaped stacking interactions with uracil justifying the relevance of W39 as highlighted by mutagenesis data. It should be noticed that W39 (TMS1) is part of a hydrophobic cleft consisting of F262 (TMS6), Y265 (TMS6), V343 (TMS8) and V189 (TMS5) contributing to the stability of the OO and Occ states where TMS1a, TMS6b, TMS8 and TMS5 are close, while in IO TMS8 and TMS5 move away as H_3_O^+^ and uracil are transported intracellularly.

## Discussion

Here we used the extensively studied at the genetic and functional level FurE protein, a eukaryotic transporter that is structurally similar to APC superfamily members, to address the mechanism of substrate/H^+^ symport using state-of-art free-energy calculations, named funnel-metadynamics (FM), focusing on the conformational rearrangements of the transporter structure that accompany transport catalysis. At variance with other binding molecular simulation methods, FM allows the sampling of the binding process without knowing *a priori* the binding mode of the ligand(s), and thus provides a unique and thorough classification of all possible binding modes. Importantly, rational mutational analysis validates the outcome and conclusions obtained via the theoretical FM calculations. Overall, this work reveals the operation mode and identifies the step-wise conformational changes that underlie the symport of uracil/H^+^ by FurE. Our principal findings are highlighted schematically in **Figure 6** and discussed in more detail below.

**Figure 6:**
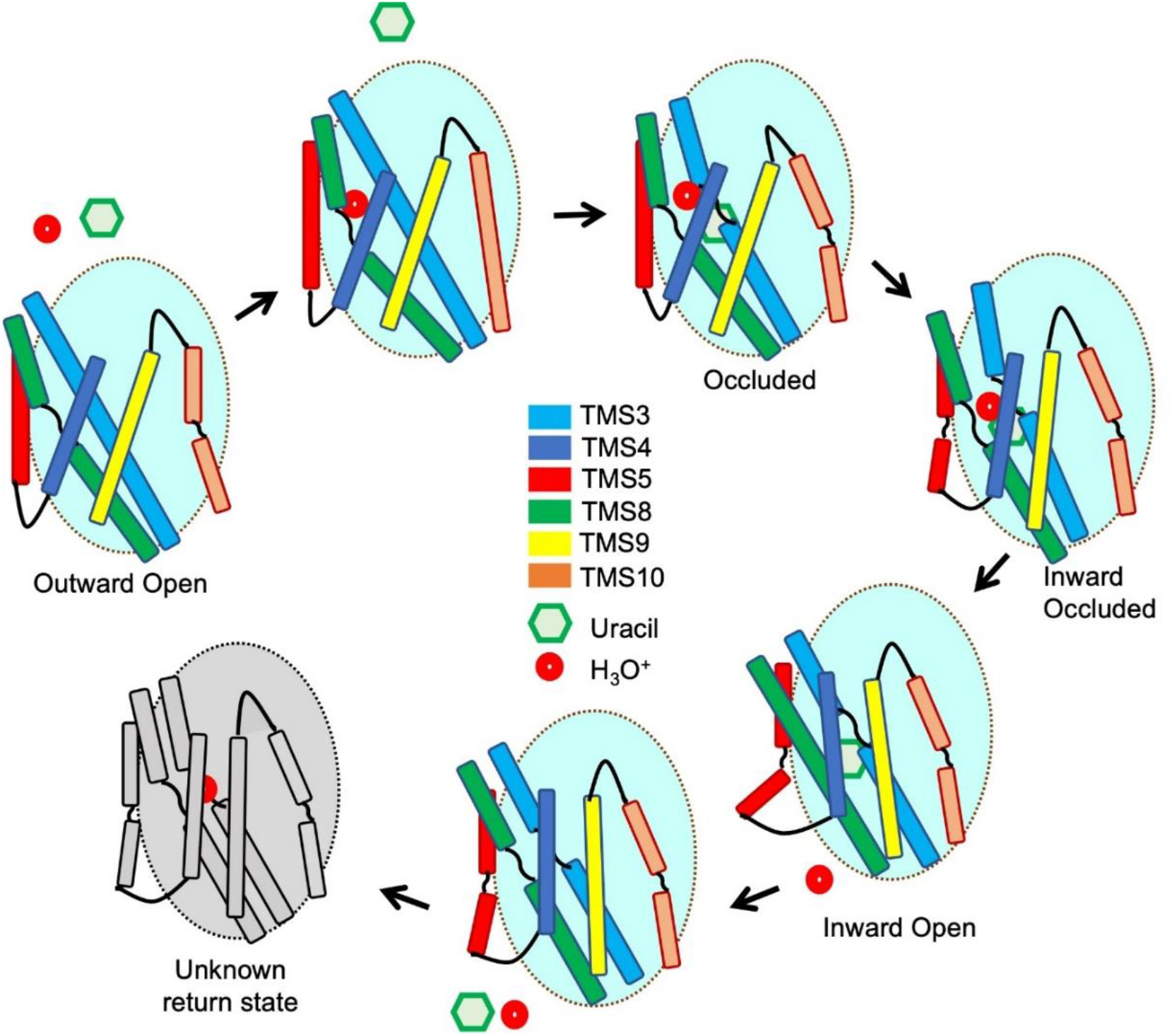
Schematic representation of the transport mechanism. The mobile FurE ‘hash’ motif helices (TMS3, TMS4, TMS8, TMS9) and outer and inner gates (TMS10 and TMS5) are shown relative to the ‘bundle’ motif, shown as cyan background, which is considered relatively immobile during uracil and H_3_O^+^ internalization. In the Outward Open (OO) state, FurE is in *apo* form. H_3_O^+^ binding results in local residue rearrangement but does not cause rearrangement of the gross tertiary structure. Uracil binding induces the closing of theTMS10 outer gate and the kink and tilt of TMS8 and TMS3, respectively, reaching the Occluded (Occ) state. H_3_O^+^ moves toward the TMS5 inner gate, which slightly bends, while TMS3 and TMS8 also display structural rearrangements, initiating the Inward Occluded (IOcc) state. After H_3_O^+^ is released in the intracellular space, TMS5 bends more, while TMS8 is not tilted anymore moving away from the ‘bundle’. TMS4 and TMS9 are shifted by TMS5, TMS8 and TMS3 bending introducing the Inward Open (IO) conformation. After the release of both H_3_O^+^ and uracil, TMS5 slightly returns to the previous bend position. An inward-facing unknown return state, probably introduced by a H_3_O^+^, is represented in grey.

A principal novelty of this work is that it addresses proton symport, by introducing H_3_O^+^ as a second distinct substrate. Thus, we obtained compelling evidence that during the whole process H_3_O^+^ interacts with three negatively charged residues, namely E51, D28 and D26. The initial binding location of H_3_O^+^ (E51) was found to be exactly at the same place where Na^+^ is co-crystallized in the homologous prokaryotic transporter Mhp1. H_3_O^+^ binding stabilized the rather flexible *apo* structure in an intermediate conformation between the initially constructed OO and Occ models (i.e. outward-occluded or OOcc). FurE-H_3_O^+^ interaction was found to trigger local amino acid rearrangements that permit Q134 to bind uracil, without promoting other major protein conformational changes, rather similar to what has been found in Mhp1.^21,35^ This local dynamic change elicited by cation-binding alone differs in other APC transporters, such as LeuT, dDAT, hDAT and SERT, where Na^+^ binding favors a fully Occ conformation.^11,12,37,38^ In FurE, only when both substrate (uracil) and H_3_O^+^ are bound, the lowest energy conformation shifted towards the Occ structure, a state where both TMS10 (outer gate) and TMS5 (inner gate) are closed. Noticeably, in the FurE Occ state we detected relatively small changes in the ‘hash’ helices. More specifically, the relative motion of W130 (TMS3) interacting with uracil elicited a small bend in the last part of TMS3 assisted by a G132, and this was followed by a similar bend of the first part of TMS8. Furthermore, TMS9 followed the movement of TMS10 and induced a small shift to TMS4 (see **Figure 4**). A network of interactions between TMS10 and TMS1b residues, namely F385, L389 and Q59, also contributed to the stabilization of the Occ state and play a critical role in the substrate specificity, as supported by the mutational analysis.

After acquiring the Occ structure, with uracil and H_3_O^+^ bound, FurE assumes an intermediate structure between Occ and IO (i.e., inward-occluded or IOcc), where H_3_O^+^ cation moved towards the intracellular domain. In this state, the N-terminal D28 loses the interaction with R264 in order to be engaged in the translocation of H_3_O^+^, while other critical rearrangements involved K199-E51-K252 and R123-D261 interactions. These events also trigger a relative motion of TMS3, TMS4 and the first part of TMS5, followed by a major tilt of TMS8 (see **Figure S8**). In this IOcc state, in which both uracil and H_3_O^+^ are still bound, we observed an initial bending of the first part of the unleashed TMS5 (at P204), which reflects the opening an inner gate. In both LeuT^39^ and DAT^40^, two sodium binding sites have been identified and related to both Occ state stabilization and substrate internalization.^10^ In FurE, K199 side chain group is located in the same position of Na ion in Mhp1 and Na2 in LeuT, and corresponds to K158 in ApcT, which is the only proton symporter crystalized today.^18,41^ In addition, the flexible side chain of K252 was very often located close to the LeuT Na1 site, in our simulations. Both K199 and K252 residues have been identified experimentally as crucial for substrate specific recognition and transport via their interaction with E51, triggering the necessary protein conformational alterations for transport activity (**Figure 4**). It thus seems that specific lys residues in H^+^ symporters might alleviate the need for Na binding needed in other APC carriers.

From the IOcc, in order to reach the IO state from, our simulations showed that H_3_O^+^ must be released first, as only in this case the FES is shifted to IO. A similar finding has been found in DAT and LeuT.^39,42–46^ Notably, however, internalization of H_3_O^+^ was accompanied by neutralization of D28 and D26 and subsequent relocation of the cytoplasmic N-terminal segment known as LID.^29^ At this state, when uracil is ready to leave the transporter, TMS5 (the inner gate) opens, the upper part of TMS3 bends, TMS8 is not tilted anymore, while TMS4 and TMS9 are relocated following the movements of TMS5, TM8 and TMS3 (**Figure S8**). H_3_O^+^ release interrupted the E51-H_3_O^+^-T336 interaction bridging the ‘hash’ and ‘bundle’ motives. This allowed the middle part of TMS8 to shift away from TMS1, which in turn resulted in maximum TMS5 bending, creating sufficient space for uracil to be in contact with the intracellular medium and leave the binding site. This is possible only by breaking a network of interactions between residues TMS1a, TMS6b, TMS8 and the cytosolic N-terminal LID, a change also contributing to substrate specificity. Overall, proton release triggers concerted conformational bending in TMS3, TMS5 and TMS8, possible due to the presence of G132, P204 and G335, respectively. Importantly, these findings suggest a deviation from the rigid-body motion of the ‘hash’ motif, shown in Mhp1 by spin label experiments, or the rocking of the ‘bundle’ domain in other APCs.^35^

After substrate dissociation FurE is in the *apo* form, and the energy minimum structure is close to IO (**Figure 4**). This finding agrees with the evidence that Mhp1 has been crystalized also in the *apo* form conformation.^20^ However, the FES minimum in the *apo* form is wide, suggesting that the transporter might assume several alternative conformations between Occ and IO. The fact that the presence of H_3_O^+^ stabilizes a FurE state close to Occ (**Figure 4C**) prompts to suggest that H_3_O^+^ binding might be essential also for the backward transition of FurE to OO.

In conclusion, we showed that H^+^/uracil binding and transport shape the energy landscape by eliciting induced-fit conformational changes that lead to sequential movements of specific TMS principally in the ‘hash’ domain, and less so in the ‘bundle’ domain, associated also with opening and closing of outer (TMS10) and inner (TMS5) gates. Our results infer that the ‘hash’ motif helices exhibit flexibility and tilt upon substrate binding in the OO-to-Occ conformational rearrangement, while in the Occ-to-IO TMS3, TMS5 and TMS8 exhibit local substrate binding-dependent flexibility, questioning the rigid rocking-movement of either the ‘bundle’ or the ‘hash’ motif, as proposed for LeuT or Mhp1, respectively. Thus, the unified picture emerging from this work is that the FurE symporter, and probably other homologous carriers, might function as a multi-step gated pore, rather than employing dramatic changes in rigid body compact domains. Finally, this work strongly supports the importance of the cytosolic N-terminal LID sequence for completion of substrate release in the cytoplasm, as also suggested by mutational analysis.^50^

## Materials and methods

### Protein Model Construction

Model of FurE was constructed based on homology modeling using Prime 2018-4 (Schrödinger, LLC, New York, NY, 2018) on Maestro platform (Maestro, version 2018-4, Schrödinger, LLC, New York, NY, 2018). Mhp1 was used as query in the three conformations: OO(2JLN), Occ(4D1B), IO(2X79), sharing with FurE a 35% similarity, while the sequence alignment was formulated according to previous work.^29^ In order to correctly represent TMS9 in the case of IO as in 2X79 IO Mhp1 crystal structure a part of it was coil, we started with the OO FurE structure and using Targeted Molecular Dynamics in plumed-v2 software,^51^ a constant force of 500000 kj/(mol*nm^2^) was applied on the Ca atoms of the helices to create FurE in occluded and inward state. The constant force was gradually turned to zero and the system was further subjected to stabilization.

### System Setup

In order to construct the protein-ligand complex CHARMM-GUI^52^ platform was used. Each model was inserted into a heterogeneous fully hydrated bilayer 120 Å × 120 Å × 120 Å, consisting of YOPC, POPI lipids and ergosterol at a ratio of 40:40:20. The membrane embedded system was solvated with TIP3P water molecules. The solution contained neutralizing counter ions and 150 mM Na^+^ and 150 mM Cl^-^. In the case that H_3_O^+^ was present, a water molecule was replaced, and the system was neutralized with Cl^-^ counter ions. The assembled simulation system consisted of ~160,000 atoms.

### Molecular Dynamics (MD) / Metadynamics

All simulations were conducted using GROMACS software, version 2019.2.^53^ CHARMM36m^54^ force field was chosen for protein and lipids, H_3_O^+^ was provided form Bryce group^55^ while the ligand and H_3_O^+^ were prepared using Antechamber^56^ and the general Amber force field.^57^ The protein orientation into the membrane was calculated using the OPM database of the PPM server.^58^ All model systems were minimized and equilibrated to obtain stable structures. Minimization was carried out for 5,000 steps with a step size of 0.001 kJ/mol applying a steepest descent followed by a conjugate gradient algorithm, and the system was equilibrated for 20ns by gradually heating and releasing the restraints to expedite stabilization. Finally, the system proceeded to further simulations free of restraints at a constant temperature of 300K using Nose-Hoover thermostat,^59^ the pressure was kept constant at 1 bar using Parrinello-Rahman semi-isotropic pressure coupling^60^ and compressibility at 4.5e-5 bar^-1^. The van der Waals and electrostatic interactions were smoothly switched off at 1.2 nm, while long-range electrostatic interactions were calculated using the particle mesh Ewald method.^61^ All bonds were constrained using the LINCS algorithm,^62^ allowing a time-step of 2.0 fs. The trajectories were further examined for structural stability by RMSD calculation of protein Ca (up to 1.2 Å) and by visual inspection with VMD platform^63^ thus ensuring that the thermalization did not cause any structural distortion.

For metadynamics^32,33,34,64^ simulations the plumed-v2 software was used.^51^

### Funnel-Metadynamics for H_3_O^+^ cation

a) The FurE transporter used was in outward-open (OO) conformation. Since no data are available concerning the binding site of the H_3_O^+^ cation, a wide area around the equivalent one of the Na^+^ cation in Mhp1 was circumvented by the funnel cone. The cone’s starting point was T332 Ca, while the cylinder had a direction towards the extracellular waters. The funnel had a switching point between the cone and cylinder region at 4.0 nm, the amplitude of the cone was 0.27 rad, the radius of the cylinder section for the unbound region at 0.1 nm, the minimum and maximum value sampled as projection of the ligand’s center of mass (COM) along the funnel axis was at 0.25 and 4.6 nm respectively, as long as, the lowest and highest value for fps.lp used to construct the funnel-shape restraint potential was at 0.00 and 4.8 nm respectively. The value for the spring constant of the funnel-shape restraint potential was set to 5000 kj/(mol*nm^2^). As collective variable (CV) was selected the distance between the Ca atom of T332 and the center of mass of H_3_O^+^ cation. The width of the Gaussian functions was set to 0.05 nm, the height to 2 kj/mol and the deposition stride to 500 simulation steps. The rescaling factor of the Gaussian function’s height was 20 as we performed well-tempered metadynamics.

b) The study of binding/unbinding process of the H_3_O^+^ cation in the cytoplasmic solvent was initiated by using well-tempered metadynamics with the FM method on FurE in the Occ state of FurE. Uracil was included in the system, placed at the binding site, whereas an upper wall of 20000 kj/(mol*nm^2^) enforced the COM in distances lower than 0.9 nm from the Cg of N341. The constructed funnel included all the possible routes that could lead the H_3_O^+^ to the exit to the cytoplasm. The cone region started at Ca of S56. The direction of the funnel axis was cytoplasm-oriented passing through Asp348 Cb atom. The switching point between the cone and the cylinder region was at 3.6 nm, the amplitude of the cone section was set to 0.5 rad and the radius of the cylinder for the unbound region to 0.1 nm. The minimum and maximum value sampled as projection of the ligand’s COM along the funnel axis was set to 0.29 and 4.2 nm respectively, the lowest and highest value for fps.lp used to construct the funnel-shape restraint potential was set to 0.2 and 4.3 nm respectively, while, the value for the spring constant of the funnel-shape restraint potential to 7000 kj/(mol*nm^2^). As CV was selected the distance between the Ca of E51 and the center of mass of H_3_O^+^ cation. The width of the Gaussian functions was calculated at 0.01 nm, the height was arranged at 2 kj/mol with a rescaling factor of the Gaussian function at 20 and the deposition stride was set to 500 simulation steps.

### Funnel-Metadynamics for Uracil

a) FM were performed aiming to highlight the binding mode of uracil in the binding site and the binding mechanism as it approaches the binding pocket from the extracellular. FurE was in the occluded state and the H_3_O^+^ cation was included in the system, as in crystallographic results of other transporters, in particular Mhp1, ion and substrate co-exist in the Occ state. In detail, the funnel used, oriented from Ca atom of V323 deep in the binding area, with direction of the funnel axis to the extracellular solute. The switching point between the cone and cylinder region was set to 2.7 nm, the amplitude of the cone section to 0.37 rad, the radius of the cylinder section for the unbound region to 0.1 nm, the minimum and maximum value sampled as projection of the ligand’s COM along the funnel axis to 0.2 and 3.3 nm respectively, the lowest and highest value for fps.lp used to construct the funnel-shape restraint potential to 0.05 and 3.6 nm respectively. The value for the spring constant of the funnel-shape restraint potential was 30000 kj/(mol*nm^2^). As CV was selected the distance between the Ca of N341 and the center of mass of uracil. The width of the Gaussian functions was 0.01 nm, the height was arranged at 2 kj/mol and the deposition stride at 500 simulation steps. The rescaling factor of the Gaussian function’s height was 20.

b) The uracil internalization process was implemented using again, well-tempered metadynamics with the FM method, on FurE transporter in IO conformation containing uracil and not H_3_O^+^, as the latter was already proved from PCV simulations that leaves first the transporter in order to allow uracil to exit too (see Main Text). The funnel was constructed as to include all the possible exiting pathways from the binding pocket to the TMS5 outer gate. The cone restraint started at backbone C atom of PF53, while the direction of the funnel axis was cytoplasm-oriented passing through S342 O atom. The switching point between the cone and the cylinder region was set to 3.4 nm, the amplitude of the cone section to 0.49 rad and the radius of the cylinder for the unbound region to 0.1 nm. The minimum and maximum value sampled as projection of the ligand’s COM along the funnel axis was set to 0.21 and 4.1 nm respectively, the lowest and highest value for fps.lp used to construct the funnel-shape restraint potential was set to 0.1 and 4.2 nm respectively, while, the value for the spring constant of the funnel-shape restraint potential was set to 30000 kj/(mol*nm^2^). As CV was chosen the distance between the backbone of A50 and the center of mass of uracil. The width of the Gaussian functions was calculated at 0.01 nm, the height was arranged at 2 kj/mol with a rescaling factor of the Gaussian function at 20 and the deposition stride was set to 500 simulation steps.

### Metadynamics Simulations with Path Collective Variable (PCV)

a) OO-to-Occ path: In this case we used the Cα atoms of the residues belonging to FurE helices involved in hash and bundle motif. This choice was found to be appropriate because the calculated FESs were well reproducible. The initial path was obtained through a carefully chosen set of frames with equally distant RMSDs, derived from a steered MD simulation where the OO FurE was biased to Occ conformation using a stable force on Ca atoms of helices. 6 frames were used to construct the path in total, while the average distance between adjacent frames was 0.13 nm. The RMSD matrix was constructed and plotted, confirming that the frames where appropriate for the calculation. The λ value calculated for s was equal to 200 nm^2^. The width of the Gaussian functions for hills deposition was 0.035 nm^2^ based on the structure fluctuation in unbiased MD, the height was arranged at 0.5 kj/mol and the deposition stride at 500 simulation steps. An upper wall of 500000 kj/(mol*nm^2^) was set to constrain the distance from the path at a value lower than 0.06, based in unbiased MD simulations of more than 200 ns where the cv’s fluctuation did not reach values higher than 0.03. If uracil is part of the system, it is constrained in the previously calculated position in the binding site with a distance restraint of 20000 kj/(mol*nm^2^) at 0.7 nm between the center of mass of the substrate and Cd atom of Q134. The same constraint was applied on the distance of H_3_O^+^ cation from Cd atom of E51 at 0.45 nm.

b) The same rationale and method were used in the Occ-to-IO case. Here, the λ value for s was equal to 110 nm^-2^, the width of the Gaussian functions for hills deposition was 0.037 nm^2^, the upper wall of 500000 kj/(mol*nm^2^) was set to constrain the z at a value lower than 0.1.

### Media, strains and growth conditions

Standard complete (CM) and minimal media (MM) for *A. nidulans* growth were used, supplemented with necessary auxotrophies at concentrations given in http://www.fgsc.net. Glucose 1% (w/v) was used as carbon source. 10 mM of sodium nitrate (NO3^-^) or 0.5 mM of uric acid (UA), xanthine (XAN) or allantoin (ALL) were used as nitrogen sources. The uracil toxic analog 5-FU was used at 100 μM in the presence of 10 mM NO3^-^ as N source. All media and chemical reagents were obtained from Sigma-Aldrich (Life Science Chemilab SA, Hellas) or AppliChem (Bioline Scientific SA, Hellas).

A *ΔfurD::riboBΔfurA::riboBΔfcyB::argBΔazgAΔ uapAΔ uapC::AfpyrGΔ cntA::riboB pabaA1 pantoB100* mutant strain named Δ7, was the recipient strain in transformations with plasmids carrying FurE mutant versions, based on complementation of the pantothenic acid auxotrophy *pantoB100*.^65^ The Δ7 strain has an intact endogenous FurE gene transporter, but this is very little expressed under standard conditions and thus does not contribute to detectable transport of its physiological substrates (UA, ALL) or to sensitivity in 5-FU^26^*A. nidulans protoplast isolation and transformation was performed as previously described*.^66^ Growth tests were performed at 37 °C for 48 h, at pH 6.8.

### Standard molecular biology manipulations and plasmid construction

Genomic DNA extraction from *A. nidulans* was performed as described in FGSC (http://www.fgsc.net). Plasmids, prepared in *Escherichia coli*, and DNA restriction or PCR fragments were purified from agarose 1% gels with the Nucleospin Plasmid Kit or Nucleospin Extract II kit, according to the manufacturer’s instructions (Macherey–Nagel, Lab Supplies Scientific SA, Hellas). Standard PCR reactions were performed using KAPATaq DNA polymerase (Kapa Biosystems). PCR products used for cloning, sequencing and re-introduction by transformation in *A. nidulans* were amplified by a high-fidelity KAPA HiFi HotStart Ready Mix (Kapa Biosystems) polymerase. DNA sequences were determined by VBC-Genomics (Vienna, Austria). Site-directed mutagenesis was carried out according to the instructions accompanying the Quik-Change^®^ Site-Directed Mutagenesis Kit (Agilent Technologies, Stratagene). The principal vector used for most *A. nidulans* mutants is a modified pGEM-T-easy vector carrying a version of the *gpdA* promoter, the *trpC* 3’ termination region and the *panB* selection marker.^26^ Mutations in FurE were constructed by oligonucleotide-directed mutagenesis or appropriate forward and reverse primers. Transformants with intact FurE alleles were identified by PCR analysis.

### Epifluorescence microscopy

Samples for standard epifluorescence microscopy were prepared as previously described.^67^

In brief, sterile 35-mm l-dishes with a glass bottom (Ibidi, Germany) containing liquid MM supplemented with NaNO_3_ and 0.1% glucose were inoculated from a spore solution and incubated for 18 h at 25 °C. The images were obtained using an inverted Zeiss Axio Observer Z1 equipped with an Axio Cam HR R3 camera. Image processing and contrast adjustment were made using the ZEN 2012 software while further processing of the TIFF files was made using Adobe Photoshop CS3 software for brightness adjustment, rotation, alignment and annotation.

### Uptake assays

FurE transport activity was measured by estimating uptake rates of [^3^H]-uracil (40 Ci mmol^-1^, Moravek Biochemicals, CA, USA), as previously described.^65^

In brief, [^3^H]-uracil uptake was assayed in *A. nidulans* conidiospores germinating for 4 h at 37 °C, at 140 rpm, in liquid MM (pH 6.8). Initial velocities were measured on 10^7^ conidiospores/100 μL incubated with A concentration of 0.2μM of [^3^H]-uracil at 37 °C. All transport assays were carried out in triplicates in at least two independent experiments. Results were analyzed using the GraphPad Prism software. Standard deviation was less than 20% in all calculations.

## Acknowledgments

We would like to thank Dr. Simone Aureli for his assistance on the computational part, Dr. Paolo Conflitti for many helpful discussions.

## Author contributions

Conceptualization: GD and EM.

Methodology and supervision:

IZ carried out the calculations with help by SR and supervision by EM and VL.

GL has constructed the initial models together with IZ.

GFP and YP carried out the experiments supervised by GD.

Writing—original draft: IZ, EM and GD

Writing—review & editing: SR, VL, GFP

## Data and materials availability

All data and materials used in the analyses are available to any researcher for purposes of reproducing or extending the analyses.

## Supplementary Materials

Supporting material file contains Table S1: The simulation time of each case of the PCV Metadynamics simulations. Figures S1-S8: Alignment of FurE and Mhp1, Details of interactions between residues in FurE models, Relative orientation of transmembrane helices of the ‘hash’ motif compared to the ‘bundle’, Funnel dimensions used for the four cases of FM simulations Comparison of the binding mode of substrates in FurE and Mhp1, RMSD diagrams, Substrate-residue interactions in different intermediate conformations and conformational changes of ‘hash’ motif helices during Occ to IO transition.

