## Supplemental Material for "Uracil/H^+^ symport by the FurE transporter challenges the rocking-bundle mechanism of transport in APC transporters"

### SUPPLEMENTARY INFORMATION

In Table S1: The simulation time of each case of the PCV Metadynamics simulations. In Figures S1-S8: Alignment of FurE and Mhp1, Details of interactions between residues in FurE models, Relative orientation of transmembrane helices of the "hash motif" compared to the "bundle", Funnel dimensions used for the four cases of FM simulations Comparison of the binding mode of substrates in FurE and Mhp1, RMSD diagrams, Substrate-residue interactions in different intermediate conformations and conformational changes of "hash" motif helices during Occ to IO transition.

**Table S1**: The simulation time of each case of the PCV Metadynamics simulations.

| **Op-to-Occc** | *apo* | $H_3O^+$ | $H_3O^+$ +uracil | uracil |
|---|---|---|---|---|
| Time(ns) | 352 | 550 | 600 | 430 |
| **Occ-to-In** | *apo* | $H_3O^+$ | $H_3O^+$ +uracil | uracil |
| Time(ns) | 330 | 390 | 314 | 298 |

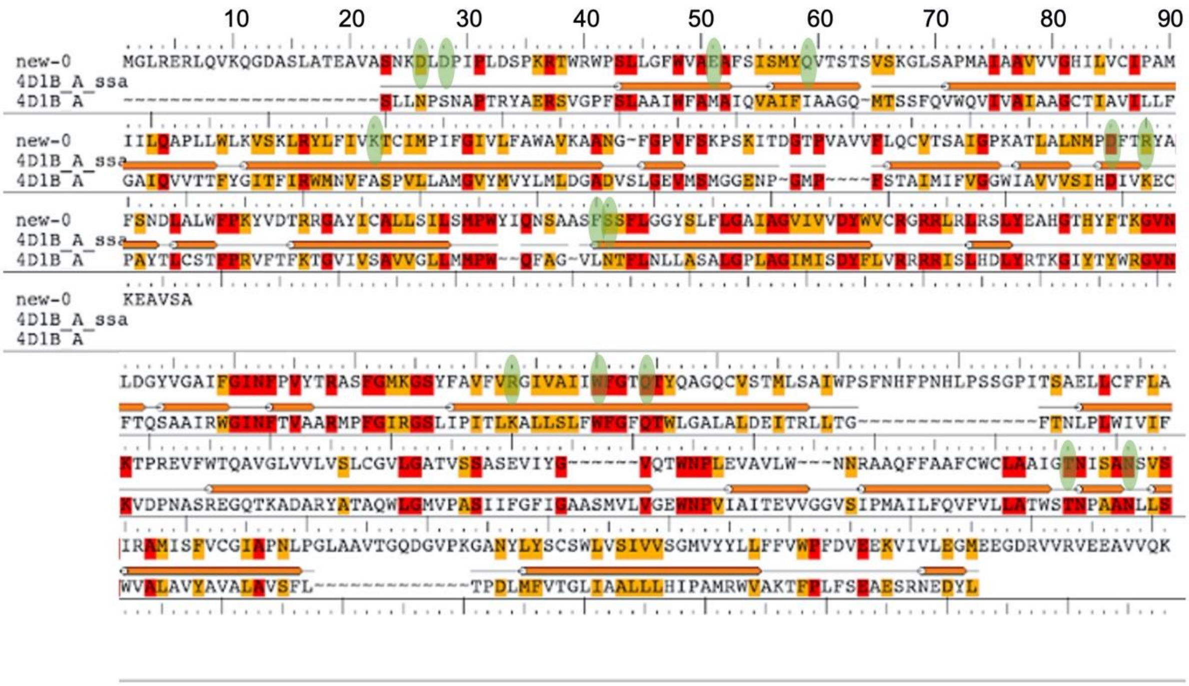


**Figure S1**: Alignment of FurE and Mhp1 (PDB ID 4D1B). Important residues studied in this work are highlighted in green.

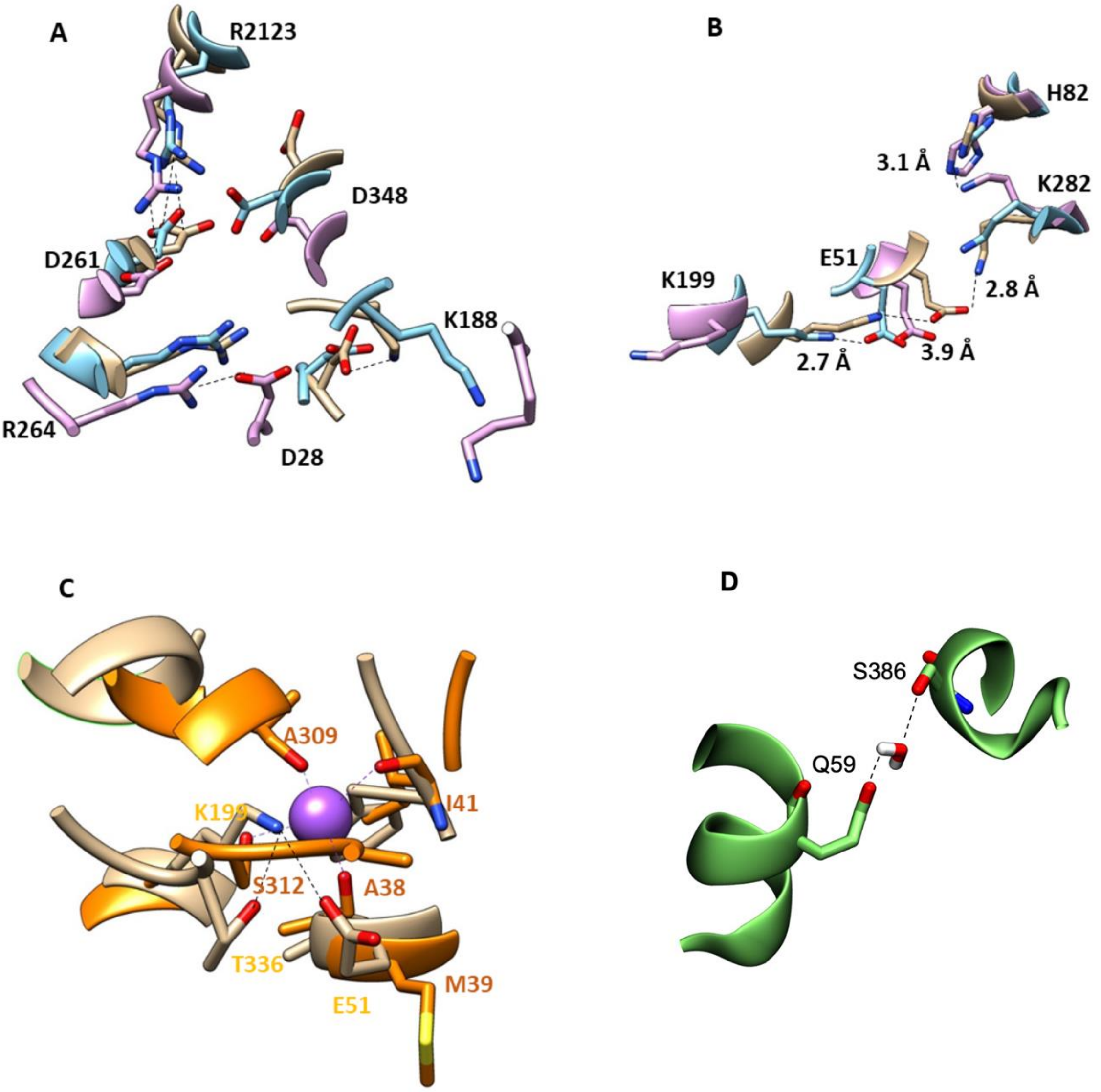


**Figure S2**. Details of interactions between residues in FurE models. (**A**) Interactions between D261-R123 (and vicinity with D348), R264-D28 and D28-K188 shown with black dashed lines and compared in the three FurE conformers (color code: OO in chaki, Occ in cyan, IO in pink). (**B)** Interaction between K199 and E51 in the different FurE conformers are shown in black dashed lines along with distances in Å (color code as in A). **(C)** In Outward Open (OO) conformation the interaction between K199-E51-T336 in FurE model (in chaki) is shown with black dashed lines. When compared with the corresponding cavity of Mhp1 (superimposed in orange), the FurE K199 amino group is located in the same position as the co-crystalized sodium (in magenta) in Mhp1 structure. Mhp1 sodium is coordinated by the S312 and T311 side chains along with A38, I41 and A309 backbone oxygens (magenta dashed lines). (**D**) Interaction between Q59 and S386 through a water molecule in FurE model MD simulation.

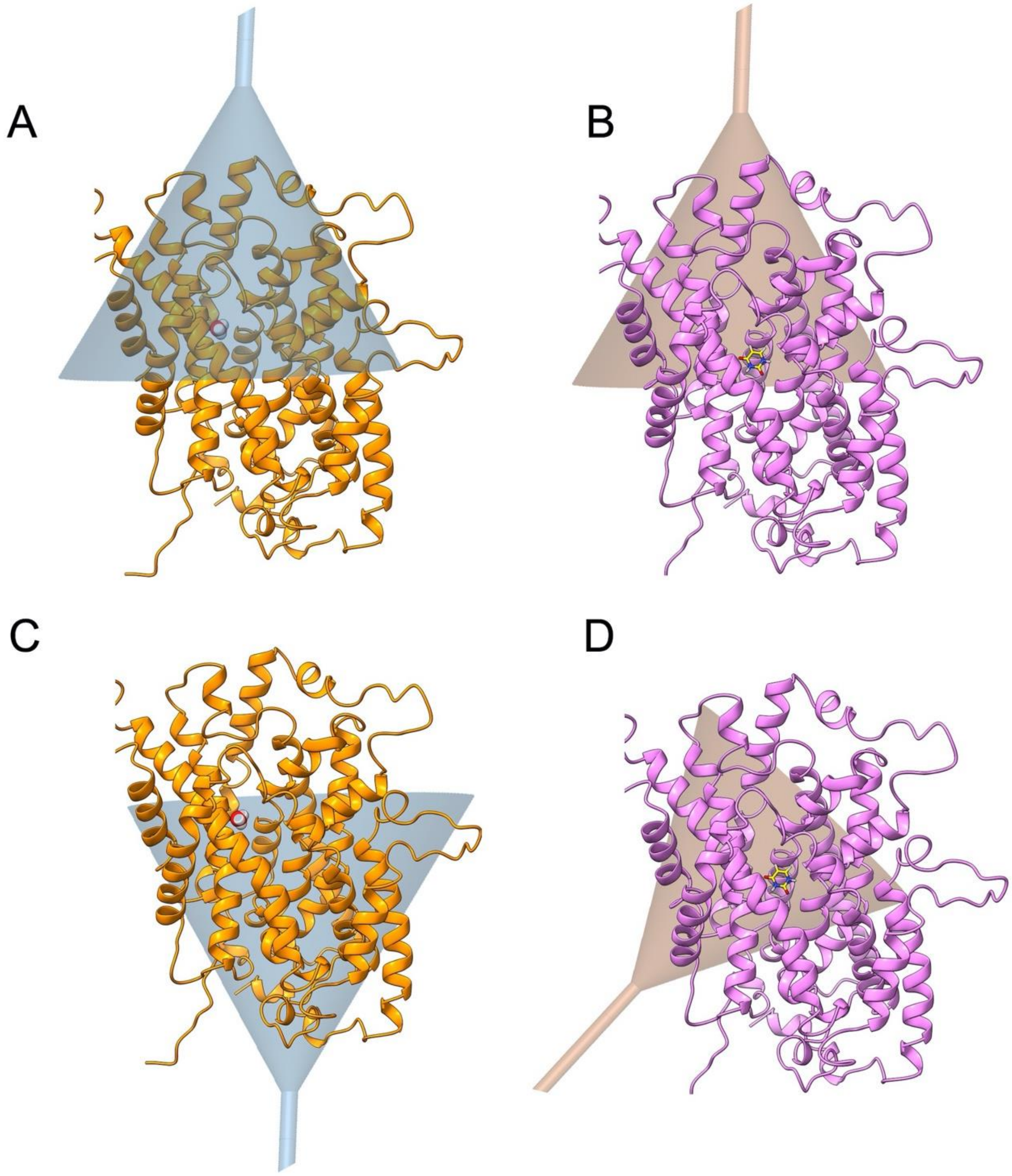


**Figure S3:** Funnel dimensions used for the four cases of FM simulations: (**A**) the funnel investigating the entrance of $H_3O^+$ in FurE transporter. (**B**) The funnel investigating the entrance of uracil in FurE transporter. (**C**) The funnel investigating the internalization process of $H_3O^+$in FurE transporter. (**D**) The funnel investigating the internalization process of uracil in FurE transporter.

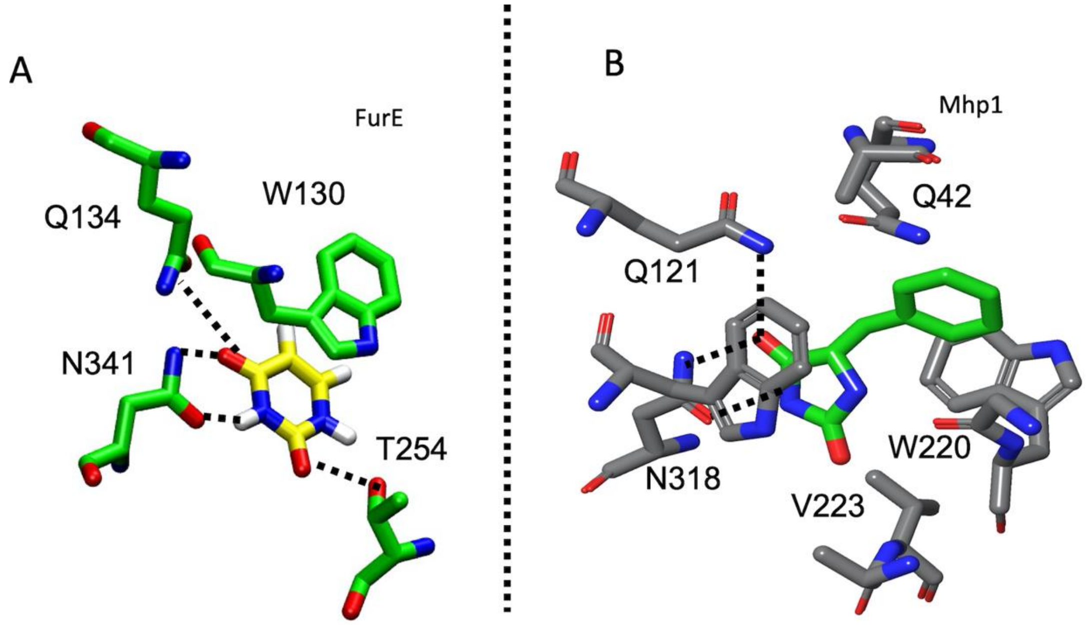


**Figure S4**: Comparison of the binding mode of (**A**) uracil in FurE with (**B**) (5S)-5-benzylimidazolidine-2,4-dione in Mhp1.

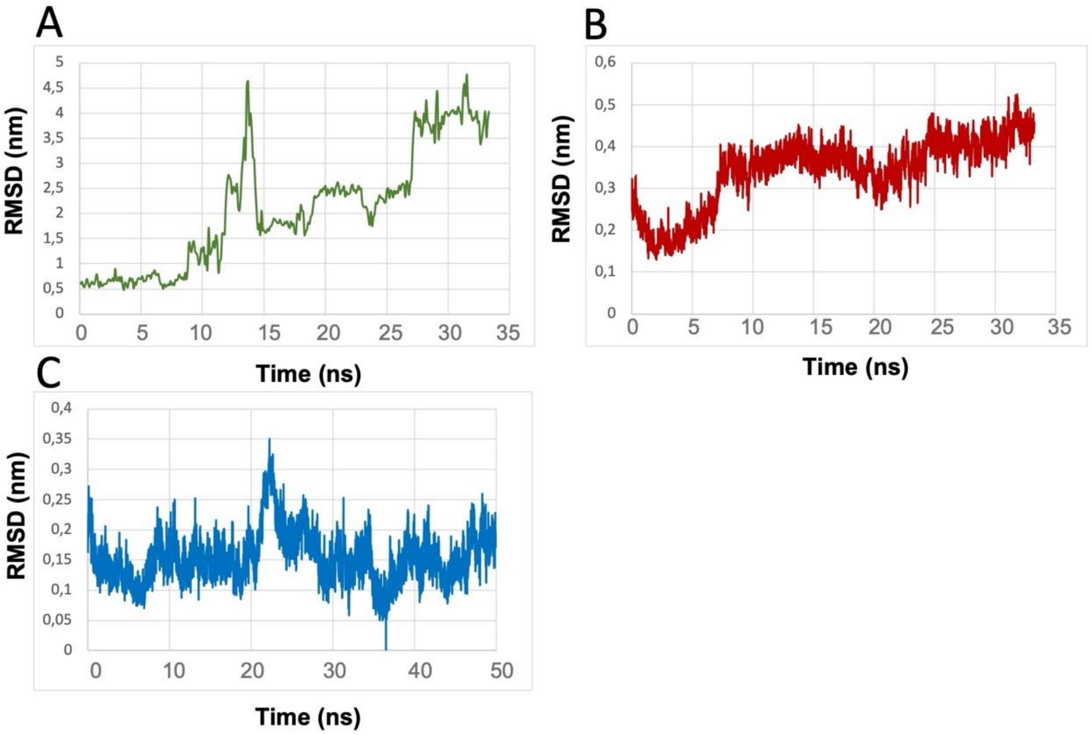


**Figure S5**: RMSD diagrams. (**A**) The RMSD fluctuation of uracil during an unbiased Molecular Dynamics simulation in the occluded FurE structure until it is out of the transporter, with uracil included and $H_3O^+$ not included. (**B**) The RMSD fluctuation of TMS10 during an unbiased Molecular Dynamics simulation in the occluded FurE structure, with uracil included and $H_3O^+$ not included. (**C**) The RMSD fluctuation of TMS10 during an unbiased Molecular Dynamics simulation in the outward *apo* FurE structure.

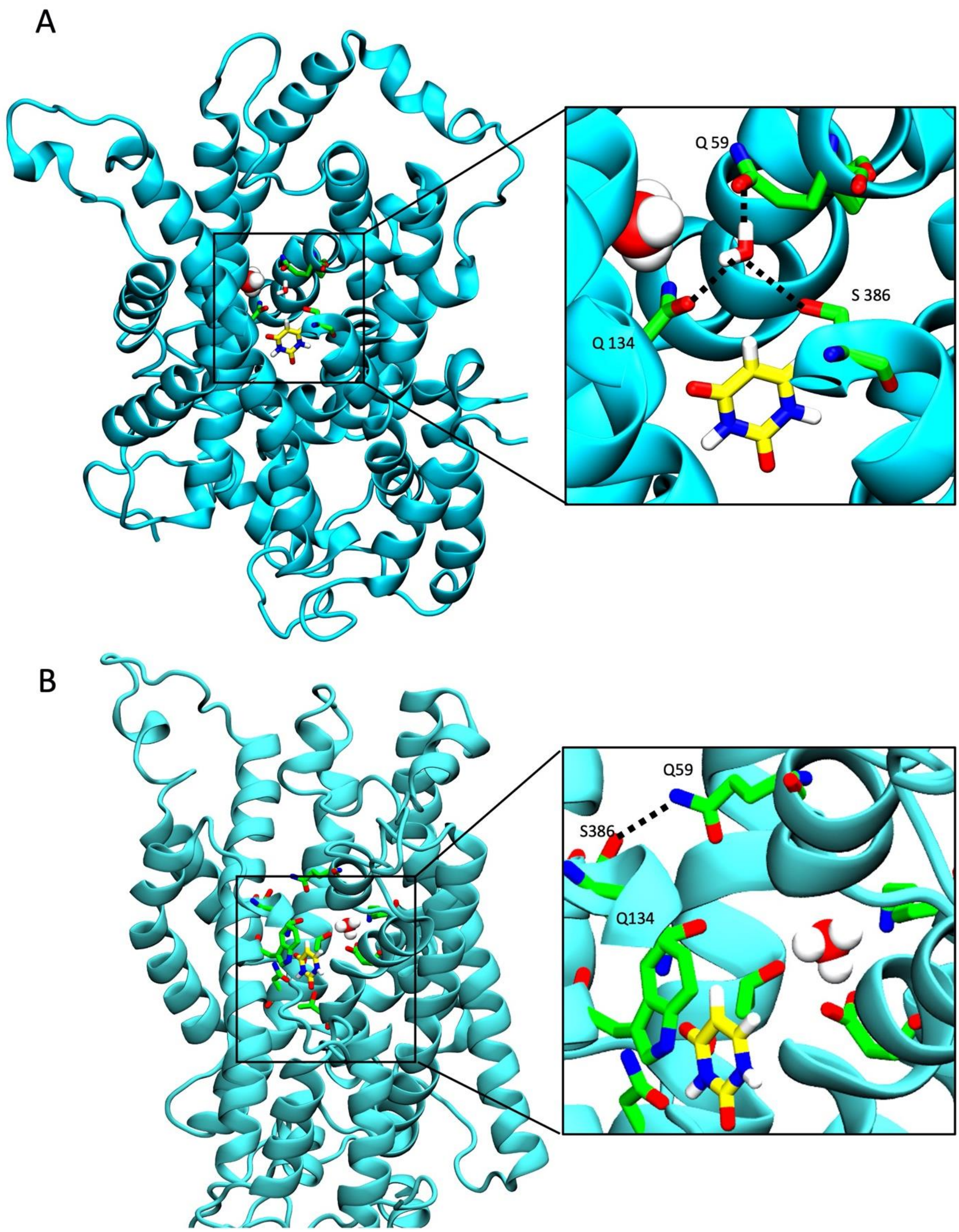


**Figure S6**: S386-Q59 interaction stabilize the Occ conformation of FurE by closing the TMS10 outer gate. (**A**) S386 interacts with Q59 through an H-bond network which involves a water molecule. (**B**) S386 forms a direct H-bond with Q59.

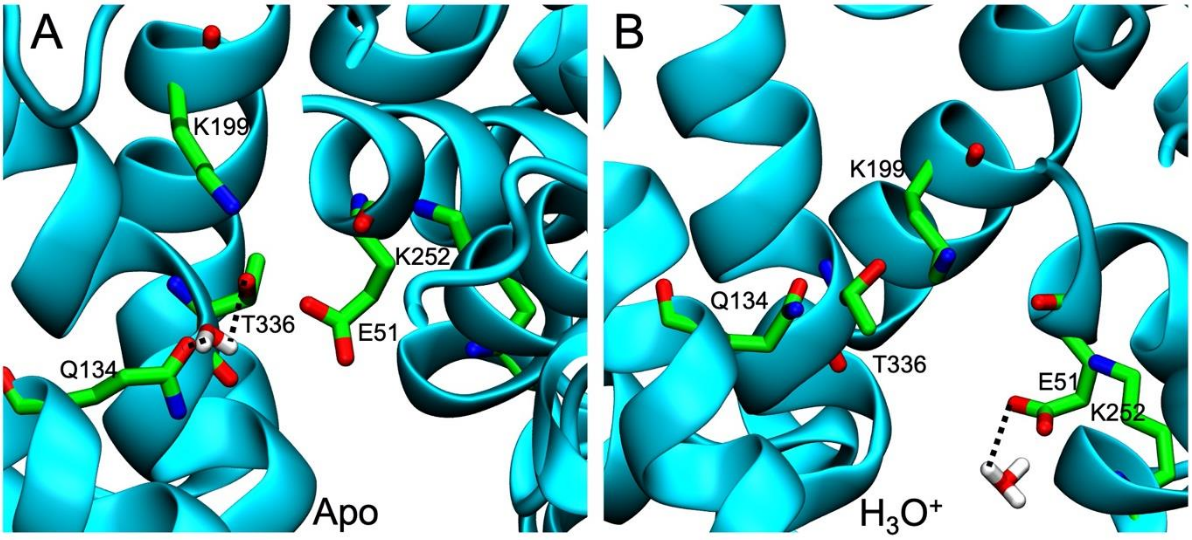


**Figure S7**: Different residue interactions in the absence and presence of $H_3O^+$ cation in the OO FurE conformation. (**A**) When FurE is in APO Form, E51 interacts strongly with K199 and less often with K252. The orientation of T336 oxygen is towards TMS3 as T336 interacts with Q134 through H-bond network involving a water molecule. (**B**) When $H_3O^+$ is present the electrostatic bond between K199 and E51 is less likely to happen, while is more likely for K252 to interact with E51. T336 side chain has also rotated by 180 degrees as it interacts with $H_3O^+$ as displayed in Figure 1B. When $H_3O^+$ is present T336 side chain oxygen is directed towards TMS5, thus the interaction with Q134 is broken and the latter is free to bind the substrate (uracil).

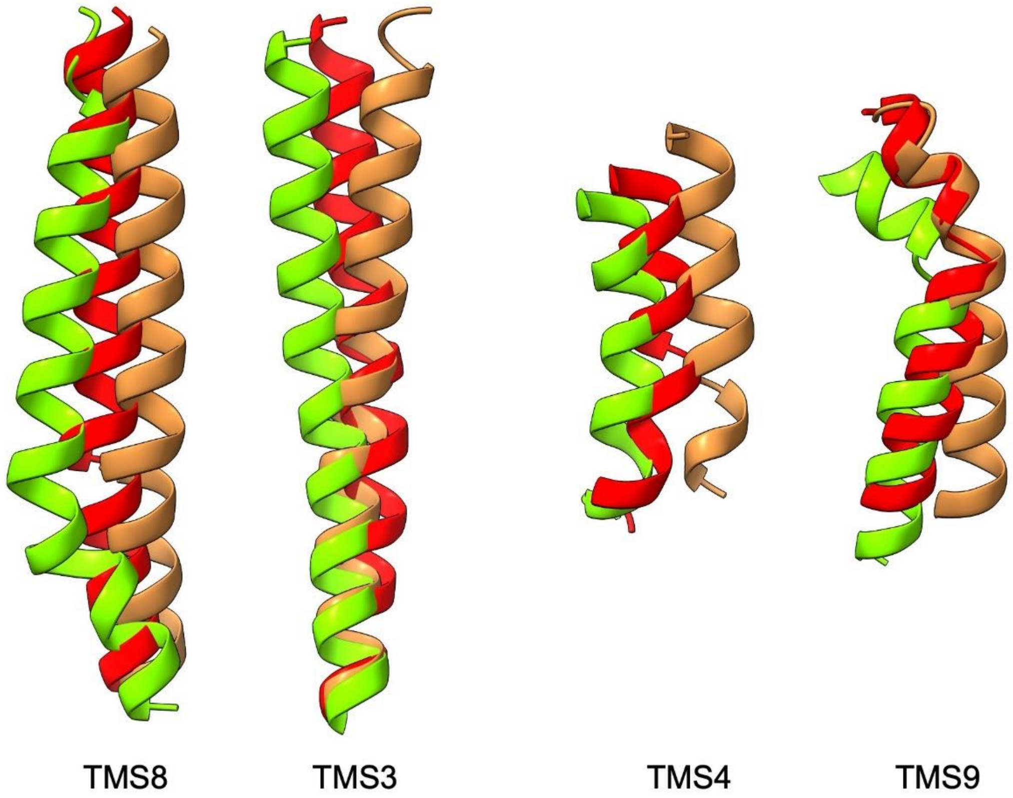


**Figure S8**: Conformational alterations of the "hash" motif helices during the Occ to IO transition
